# Assessing the Validity of Leucine Zipper Constructs Predicted in AlphaFold2

**DOI:** 10.1101/2024.10.14.618350

**Authors:** Isobel Mitic, Katharine A. Michie, David A. Jacques

## Abstract

AP-1 transcription factors are a network of cellular regulators, that combine in different dimer pairs to control a range of pathways involved in differentiation, growth, and cell death. They dimerise via leucine zipper coiled-coil domains, that are preceded by a basic DNA binding domain. Depending on which AP-1 transcription factors dimerise, different DNA sequences will be recognised resulting in differential gene expression. The affinity of AP-1 transcription factors for each other dictates which dimers form. The relative concentration of AP-1 transcription factors varies with tissue type and environment, adding another layer of control to this integral network of cellular regulation. The development of artificial intelligence (AI) protein structure prediction programs gives us a new technique to investigate or predict how dimerization effects combinatorial control. AlphaFold2 and AlphaFold-Multimer are AI programs that predict 3D structures of proteins using primary sequence as their only input, even if there is no homologous model available. To fully realise the potential of AI for structural biology, it is essential to understand its current capabilities and limitations. In this study we used the classical example of an AP-1 dimer: Fos and Jun, to interrogate how AlphaFold2 and AlphaFold-Multimer model leucine zipper domains, and if AlphaFold-Multimer can be used to differentiate between probable and improbable dimer interfaces. We found that AlphaFold-Multimer predicts highly confident leucine zipper dimers, even for dimer pairs, such as the FosB homodimer, for which electrostatics are known to prevent their formation in vivo. This is an important case study concerning high-confidence, but low-accuracy protein structure prediction.

**statement:** Artificial intelligence (AI) programs that predict protein structures, like AlphaFold, could transform structural biology by speeding up the experimental process. However, it is important to grasp the capabilities and limitations of these AI tools. This study examines how AlphaFold identifies structural features, specifically a leucine zipper, while not considering other factors like electrostatic interactions, using the well-studied transcription factors Fos and Jun as a case study.

## Introduction

Fos and Jun were first identified as the oncogenic viral protein vFos and vJun, isolated from avian sarcoma virus, before their eukaryotic homologs were cloned and characterised (Vogt, Bos and Doolittlet, 1987). Since then, Fos and Jun have been incorporated into the larger category of Activator Protein 1 (AP-1) transcription factors, which are a ubiquitous family of homo or heterodimers with a coiled-coil Leucine zipper (L-zip) dimer interface. (Yin *et al*., 2017; Bejjani *et al*., 2019) (See Figure *1*. Panel A)

**Figure 1:**
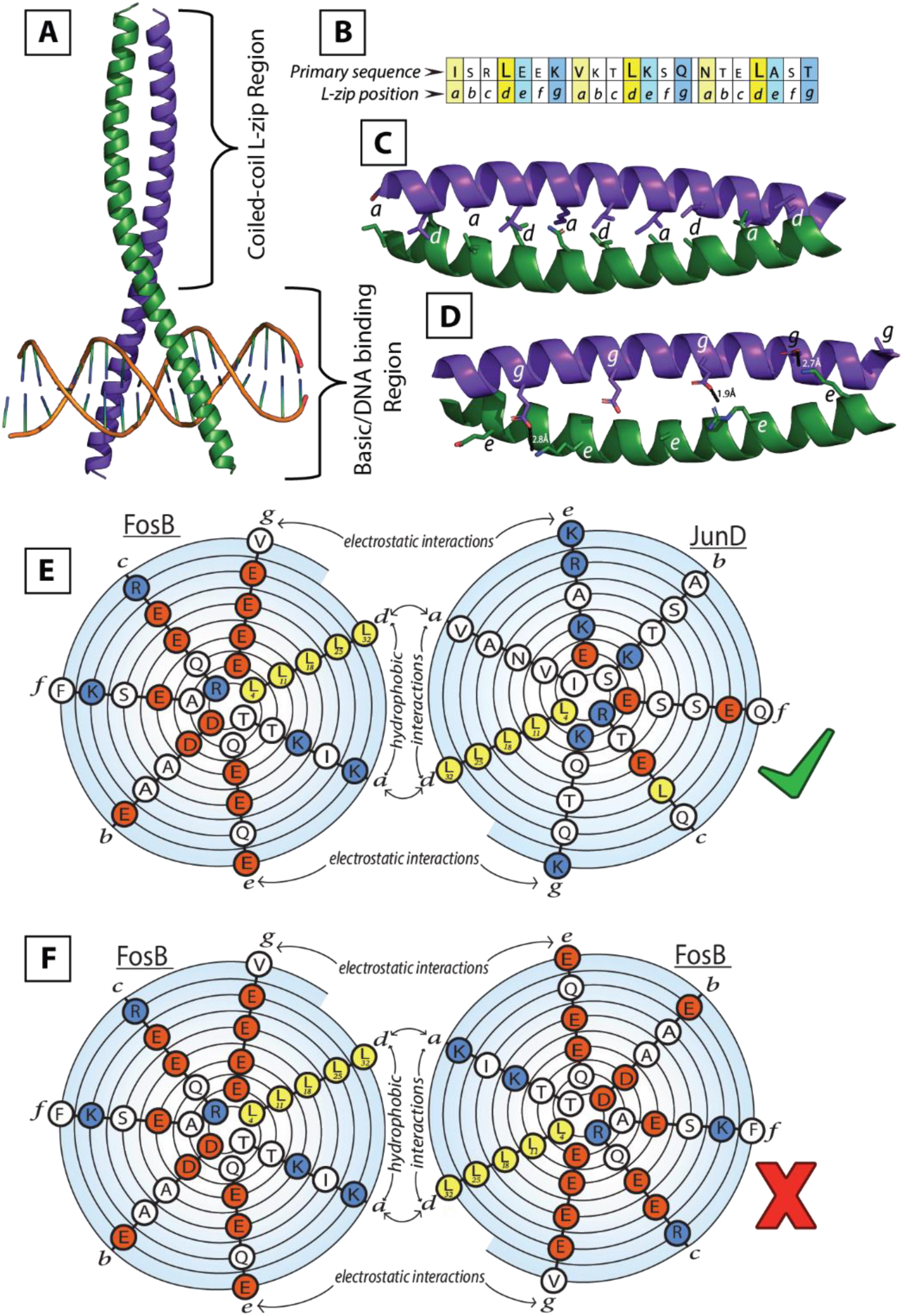
The structure of a Leucine-zipper. FosB is shown in purple, JunD is shown in green. A: FosB and JunD dimer bound to DNA. B: Primary sequence of three heptad repeats from JunD, a and d in yellow, e in cyan, g in blue. C: L-zip region of FosB-JunD dimer, a and d residues shown as sticks. D: L-zip region of FosB-JunD dimer, g residues from FosB and the e residues from JunD shown as sticks. Interchain bonds shown as dashed black lines. E: Spiral diagram of the primary sequence of FosB and JunD. The N-terminus of the L-zip starts in the centre of each spiral, and seven residues are shown over two turns of the spiral diagram. Each spiral is orientated to mimic the structure of the coiled-coil L-zip. Leu residues are shown in yellow, negatively charged residues are shown in red and positively charged residues are shown in blue (Branden and Tooze, 1991) F: Spiral diagram of the primary sequence of FosB in a homodimer.

AP-1 transcription factors, including Jun, Fos, Maf and ATF proteins, regulate a wide range of biological pathways through combinatorial control. These pathways include cell proliferation and apoptotic signalling, classifying AP-1 transcription factors as both oncogenic and tumour suppressors (Eferl and Wagner, 2003). Furthermore, cell signalling pathways controlled by AP-1 transcription factors are frequently disrupted or dysregulated by viral and intracellular bacterial infections, contributing to the pathogenicity of these infectious agents (Zachos, Clements and Conner, 1999; Gazon *et al*., 2012; Krämer *et al*., 2015). The expression levels of each individual AP-1 protein vary depending on cell type and environment, and the relative affinity that AP-1 proteins have for each other varies depending on the primary sequence of their L-zip coiled-coil domain. This forms a complex network of transcriptional regulators, based on dimer formation and affinity (Eferl and Wagner, 2003; Bejjani *et al*., 2019). Perhaps unsurprisingly, the AP-1 L-zip dimer interface has been analysed in atomic detail to understand dimer affinity, specificity and formation (Vinson *et al*., 2002; Newman and Keating, 2003; Vinson, Acharya and Taparowsky, 2006).

AP-1 L-zips are formed of 4-6 heptad repeats, and each residue in the heptad is labelled *a-g*. Residue *d* is normally a leucine, and residue *a* is normally aliphatic and non-polar; together they form the hydrophobic core of the coiled coil. (See Figure *1* Panel B, C) Residues *e* and *g* are usually polar or charged residues, and they interact to stabilise, or destabilise, the dimer interface, either side of the hydrophobic core. (See Figure *1* Panel B, D) Variation within these heptad repeats dictates the ability of different monomers to dimerize (Vinson *et al*., 2002; Vinson, Acharya and Taparowsky, 2006). Fos and Jun are a canonical example of this.

The Fos and Jun families include FosB, c-Fos, JunB, c-Jun and JunD. These proteins have been used as textbook examples of coiled-coil structure motifs for decades (Branden and Tooze, 1991). It is well documented that Fos and Jun form stable heterodimers, and Jun can homodimerize, but Fos cannot form stable homodimers due to repelling charges within the L-zip dimer interface (Branden and Tooze, 1991).

As shown in Figure 1 Panel D, E, the *g* position residues of FosB are mostly glutamates, which interact with the positively charged *e* position residues of JunD. The favourable electrostatic interaction allows a stable dimer to form. Conversely, electrostatic repulsion between these same *g* positioned glutamates and those in position *e,* prevent FosB from forming stable homodimers. (See Figure *1* Panel F) (Vinson, Acharya and Taparowsky, 2006; Chen, De Jong and Shin, 2012). Crucially, some dimer pairs are possible, and others are forbidden, dictated by attractive or repulsive electrostatic interactions. Being able to accurately discriminate functionally-relevant L-zip interactions from those that are forbidden has the potential to inform our understanding of the transcriptional reprogramming that occurs in cancers and in response to certain intracellular pathogens (Eferl and Wagner, 2003; Kuhlmann *et al*., 2007; Gazon *et al*., 2012; Krämer *et al*., 2015). The ability to predict L-zip dimer structures accurately using AI would give us a new approach to analyse numerous different theoretically possible dimer interfaces.

In 2020, the AI program AlphaFold2 (Jumper *et al*., 2021) was shown to predict highly accurate protein structures in the 14th Critical Assessment of protein Structure Prediction. AlphaFold2 incorporates a neural network (Evoformer) that predicts 3D protein structures from a primary sequence, based on multiple sequence alignments. In addition to this, AlphaFold-Multimer was developed in March 2022, and uses a training set of oligomeric structures to predict the structure of multimeric proteins with known stoichiometry (Evans *et al*., 2022). In AlphaFold2 and AlphaFold-Multimer, the proximity of protein residues is represented by a graphical network which encodes positional information only. It cannot explicitly account for other effects, for example electrostatics, that can modify proximity (Jumper *et al*., 2021; Evans *et al*., 2022). We used FosB and JunD as a test case to investigate the capabilities and limitations of Alphafold2 and AlphaFold-Multimer when positional information for residues is only one factor to take into account.

## Results

### AlphaFold2 predicts FosB leucine zipper homo- and heterodimer structures with indistinguishable confidence

AlphaFold2 (Jumper *et al*., 2021) predicted the FosB monomer had an α-helix covering residues P152-G221. The rest of the protein was unstructured. For the JunD monomer, AlphaFold2 predicted two α-helices: V128-A165 and M263-S334. (See Supplementary Figure 1) It is notable that AlphaFold2 predicts these structured helices in FosB and JunD in the absence of their binding partners. To date, it is not clear if AP-1 transcription factors retain their secondary and quaternary structures in the absence of DNA *in vivo* (Yin *et al*., 2017).

Alphafold-Multimer (Evans *et al*., 2022) predicted the FosB/JunD heterodimer (See Figure 2 Panel B) contained a high confidence coiled-coil dimer interface. The predicted structures of the FosB (See Figure 2 Panel A) and JunD (See Figure 2 Panel C) homodimers had very similar structures. The coiled coil structured regions covered residues 152±1-219±1 of FosB and 263±1-334±1 of JunD. This was consistent and reproducible across all five ranked structures (See Supplementary Figure 2, 3, 4). These results are also consistent with the crystal structure of the FosB/JunD heterodimer (PDB code: 5VPF), which contains only residues 153-217 and 267-330, respectively. Upon first inspection, the Alphafold-Multimer structures of the FosB and JunD homodimers showed remarkably little difference to the FosB-JunD heterodimer.

**Figure 2:**
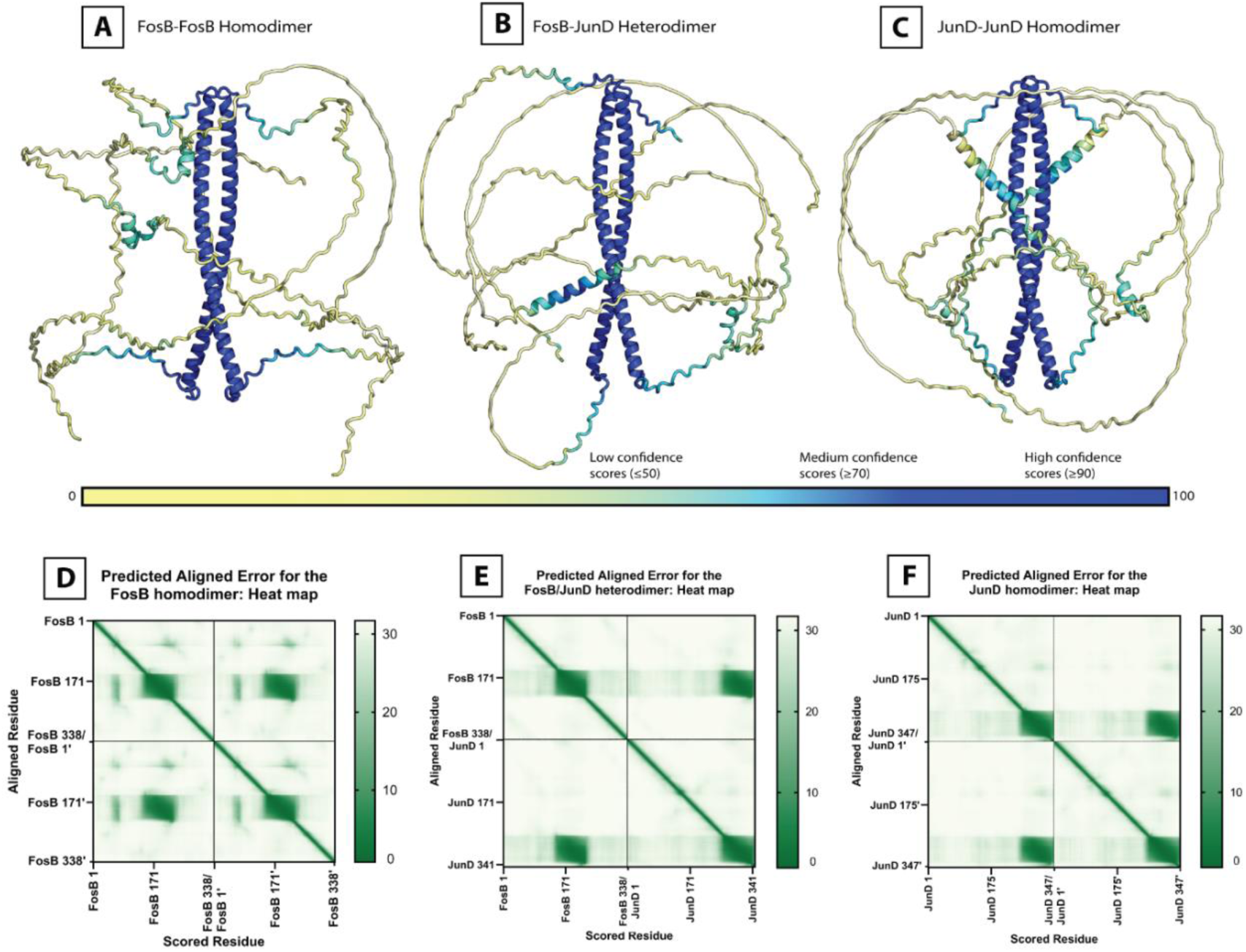
AlphaFold-Multimer predicted structures coloured by confidence score and the corresponding Predicted Alignment Error plots. A: The structure of the FosB homodimer coloured by confidence score; B: The structure of the FosB-JunD heterodimer coloured by confidence score; C: The structure of the JunD homodimer coloured by confidence score; D: The PAE scores for the FosB homodimer shown in a heatmap; E: The PAE scores for the FosB-JunD heterodimer shown in a heatmap; F: The PAE scores for the JunD homodimer shown in a heatmap.

Alignments between the crystal structure of the FosB-JunD heterodimer, and the three different AlphaFold-Multimer constructs confirmed that the coiled-coil regions are accurate representations of an L-zip. Within all three dimers, the L-zip Leu (*d*) residues and the *a* residues are positioned correctly to form a hydrophobic core between the coiled-coils. (See Supplementary Figure 5.) The RMSD for these alignments are all <1Å. (See Table 1)

**Table 1:**
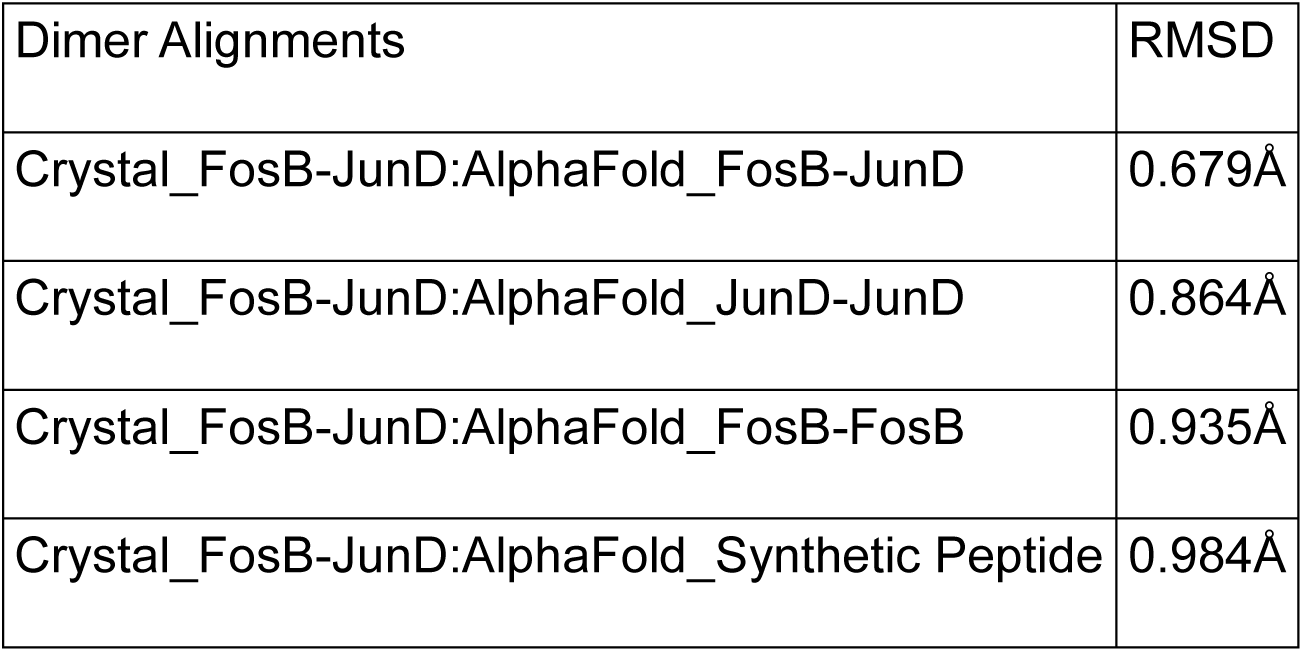
RMSD of structural alignments.

AlphaFold2 and AlphaFold-Multimer calculate a predicted Local Distance Difference Test (pLDDT) score for each residue in the predicted structure. These pLDDT scores range from 0 to 100. A high pLDDT score is indicative of accurate local geometry within a protein chain (Mariani *et al*., 2013). pLDDT scores of >90 are considered to be high confidence, and any score under 70 is considered to be low to very low confidence (Mariani *et al*., 2013; Jumper *et al*., 2021). By definition, intrinsically disordered domains will have low pLDDT scores (Mariani *et al*., 2013).

The pLDDT scores for the structured L-zip regions for the three dimer structures were all high: ≥90. The mean pLDDT score for the FosB(T180-V214)-JunD(I293-K327) heterodimer L-zip region was 97.7, for the JunD(I293-K327) homodimer it was 95.1, and for FosB(T180-V214) homodimer it was 94.4. These high pLDDT scores indicate that AlphaFold-Multimer is highly confident in the predicted structure of all three of these coiled-coil L-zip regions. Large regions of the predicted FosB and JunD dimer structures had no recognisable secondary structure and very low pLDDT scores of 20 to 40; it is reasonable to infer that these regions are intrinsically disordered based on experimental evidence (Yin, 2019; Kumar *et al*., 2022). (See Supplementary Figure 2, 3, 4)

AlphaFold-Multimer can produce an “interface predicted template modelling” (ipTM) score for each ranked structure. The ipTM scores are a global measure, designed to give the user an indication of the quality of the overall predicted structure, and have been adjusted to effectively evaluate multimeric protein complexes. In general, ipTM scores ≥0.8 indicate a high confidence predicted structure, and ipTM scores of between 0.6-0.8 indicate medium confidence (Evans *et al*., 2022). As can be seen in Figure 3 Panel A, the ipTM scores for the FosB and JunD homo/heterodimers were all low, at approximately 0.3. This was true for the dimers that are known to form *in vivo* (FosB/JunD and JunD/JunD), as well as the dimer unlikely to form *in vivo* (FosB/FosB).

**Figure 3:**
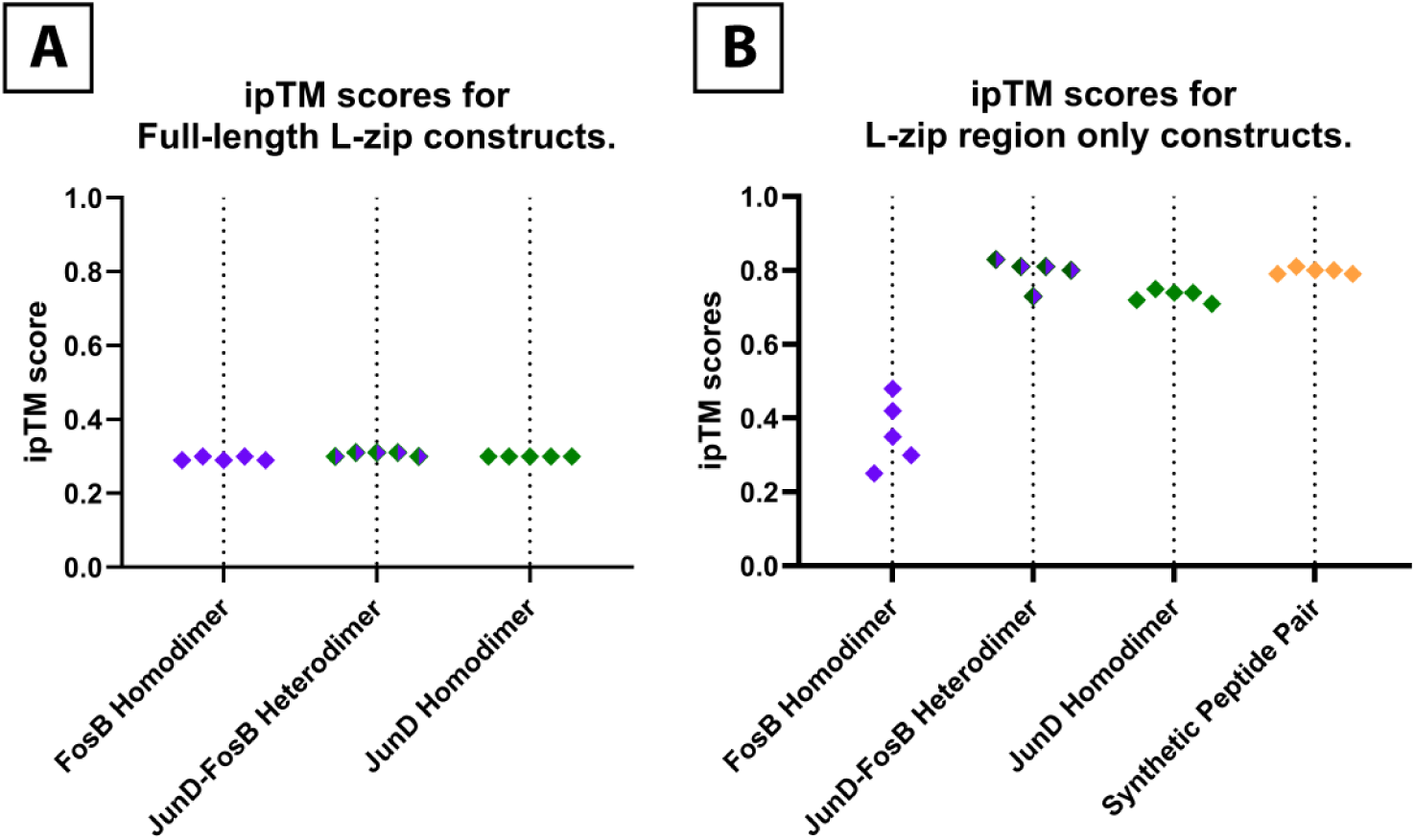
ipTM scores for the top five ranked structures for each dimer construct. Panel A: ipTM scores for the full-length FosB/JunD dimer pairs. Panel B: ipTM scores for the L-zip region only of FosB(T180-V214) and JunD(I293-K327), and the synthetic peptides pair.

We hypothesised that the low ipTM scores were due to the majority of these protein complexes being intrinsically unstructured, leading to the low pLDDT scores for these sections and lowering the overall ipTM score. To test this hypothesis, we ran three peptide pairs through AlphaFold-Multimer that represented the L-zip region of FosB and JunD. Residues T180-V214 of FosB and residues I293-K327 of JunD were run as homodimers and a heterodimer, using the exact same conditions as our full-length dimer constructs. The ipTM scores for these L-zip only regions were strikingly different to the full-length constructs. Notably, the FosB-JunD heterodimer (L-zip region only) had the highest ipTM scores, (See Figure 3 Panel B), indicating a high confidence in this dimer interface. The FosB homodimer (L-zip region only) had much lower ipTM scores of between 0.25 and 0.48. These differing ipTM scores were the only indications of AlphaFold-Multimer distinguishing between L-zip constructs that are likely or unlikely to form *in vivo*.

Predicted Alignment Error (PAE) scores are a measure of confidence of the relative position of pairs of residues within the predicted structure. Therefore, PAE scores can be used to assess the quality of the positioning of different domains within the predicted structure, or between monomers in multimeric structures (Elfmann, Org Stülke and Stülke, 2023). PAE scores are measured in Ångströms (Å), and are defined as the expected position error at residue *x* if the actual and predicted structures are aligned at residue *y.* A low PAE indicates high confidence in the relative positions of a pair of residues, and a high PAE indicates low confidence (Varadi *et al*., 2021; Evans *et al*., 2022; Elfmann, Org Stülke and Stülke, 2023).

The most common way to view and interpret PAE scores are through heatmaps like those in Figure *2*. Each residue in the full multimer structure is numbered down the x and y axis, creating a matrix of every possible pair of residues within the full structure. The diagonal line seen down the centre of each heat map shows where every residue is aligned against itself, and the PAE score is always low, by definition (Varadi *et al*., 2021; Evans *et al*., 2022; Elfmann, Org Stülke and Stülke, 2023).

As can be seen in Figure *2* panel E, the PAE score heat map for the FosB-JunD dimer contained two areas of low PAE scores for intra-chain residue pairs (these can be seen along the diagonal), and two areas of low PAE scores for inter-chain residue pairs (seen off the diagonal). The inter-chain residue pairs exclusively fell within the L-zip region of the FosB-JunD dimer. This was to be expected for a well characterised dimer like Fos and Jun. Similar areas of low PAE scores for inter-chain residue pairs could be seen for the JunD homodimer and the FosB homodimer (See Figure 2 Panel D, F). Ordinarily, areas of low PAE scores like this would indicate an accurate predicted structure (Elfmann, Org Stülke and Stülke, 2023). It was remarkable that AlphaFold-Multimer still predicted low PAE scores for the L-zip region of the FosB homodimer, when it has been shown that FosB cannot stably homodimerize *in vivo* (Newman and Keating, 2003; Vinson, Acharya and Taparowsky, 2006).

PAE scores can also be expressed as pseudo-bonds. These pseudo-bonds can be drawn between any pair of residues within the predicted structure, but for convenience are usually drawn between residues that are close together in space, and each bond has an associated PAE score. We examined the L-zip region of our three predicted dimer structures to see if there was a difference between the network and PAE scores for the pseudo-bonds connecting the *e* and *g* residues in each dimer interface. Within each predicted dimer construct, we isolated the *e* and *g* residues from each monomer and drew the pseudo-bonds between them and the *e* and *g* residues from the corresponding monomer. We only looked at pseudo-bonds between residues that were <10Å apart. It should be noted that the pseudo-bonds were drawn between the Cα of each residue, even though any atom in the residue could be <10Å from an any atom in another residue.

As can be seen in Figure 4, all the pseudo-bonds between the *e* and *g* residues of the FosB-JunD dimer had low PAE scores. The PAE scores for the pseudo-bonds in the JunD homodimer and the FosB homodimer were also all low, indicating a high-quality predicted structure. There were more longer ranging pseudo-bonds in the FosB-JunD heterodimer and the JunD-JunD homodimer, possibly because of how the sidechains for the *e* and *g* residues for these two dimers are positioned relatively close together. In the FosB homodimer, the *e* and *g* residue sidechains were positioned further apart from each other, resulting in a less extensive network of pseudo-bonds. It can be seen in the FosB homodimer structure in Figure 4, that there are pseudo-bonds with low PAE scores connecting multiple unfavourable Glu-Glu *e* to *g* pairs. These closely resemble the low scoring pseudo-bonds between favourable Lys-Glu or Arg-Glu *e* to *g* pairs seen in the FosB-JunD heterodimer. Close examination of the pseudo-bonds and their corresponding PAE scores of the three structures shows relatively little difference between the different dimer interfaces, despite the low likelihood that the FosB homodimer can form *in vivo*.

**Figure 4:**
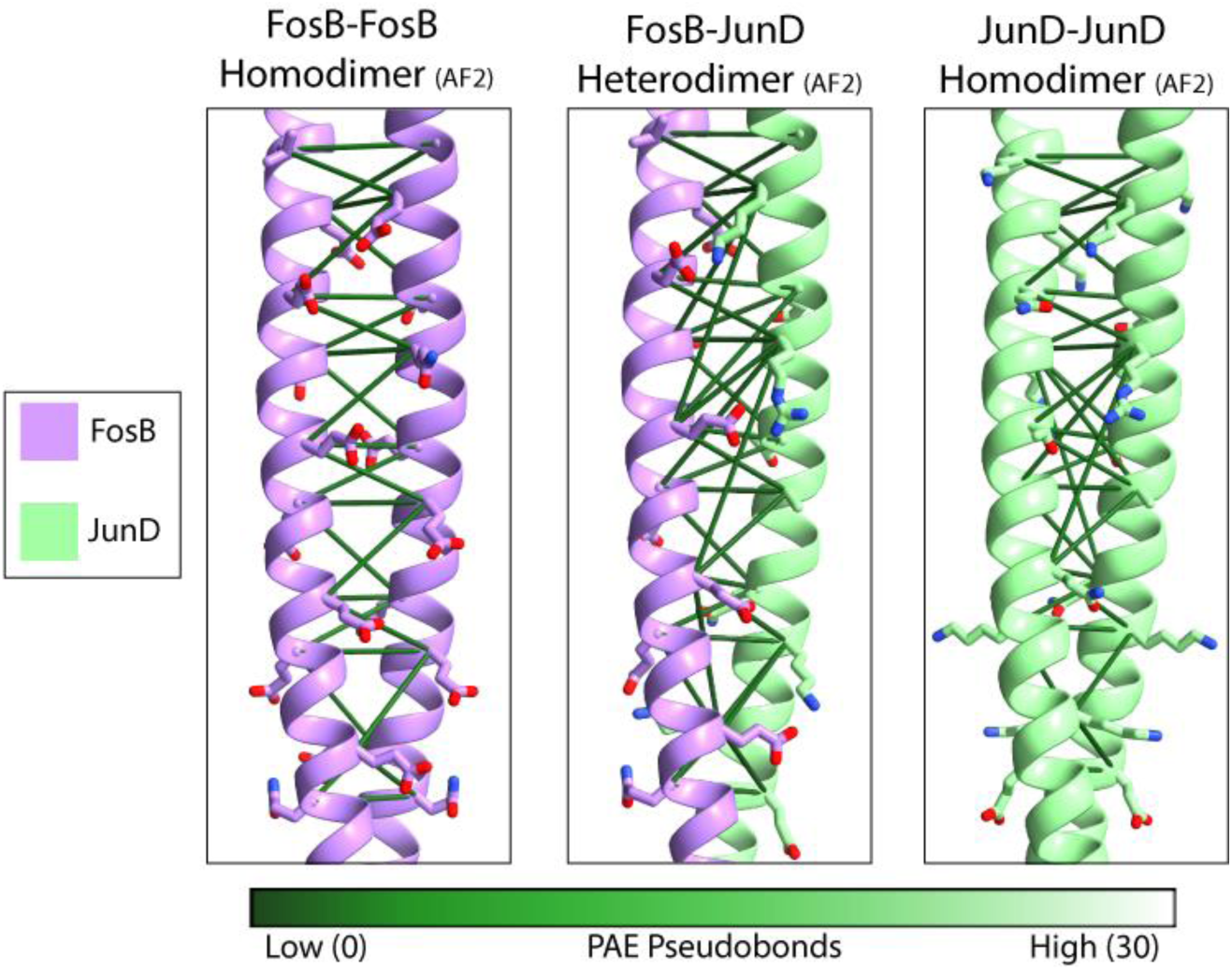
Close-ups of the L-zip region of the three different dimer structures produced in AlphaFold-Multimer. Interchain pseudo-bonds between e and g residues <10Å apart are shown and colour coded by PAE score. (Intrachain pseudo-bonds have not been shown.) Throughout: FosB shown in purple, and JunD is shown in green. The e and g positioned residues are shown as sticks and coloured by atom.

### Electrostatic surface potentials indicate a highly unlikely dimer interaction in the FosB homodimer

Lastly, we wanted to visualise the interaction between the charged sidechains in our different dimer constructs. We did this by calculating the surface potential of each individual α-helix in the L-zip constructs, and examined how each charged surface interacted with its respective partner.

The surface potential of the AlphaFold-Multimer FosB/JunD heterodimer showed that the JunD L-zip *e* position residues formed a ridge of net-positive charge along the α-helix. The negatively charged *g* residues on FosB interact with this positively charged surface favourably. (See Figure 5 Panel A.) These opposingly charged surface potentials recapitulated what was seen in the crystal structure of the FosB-JunD heterodimer (See Figure 5 Panel B): the two helices forming opposingly charged ridges that favourably interact.

**Figure 5:**
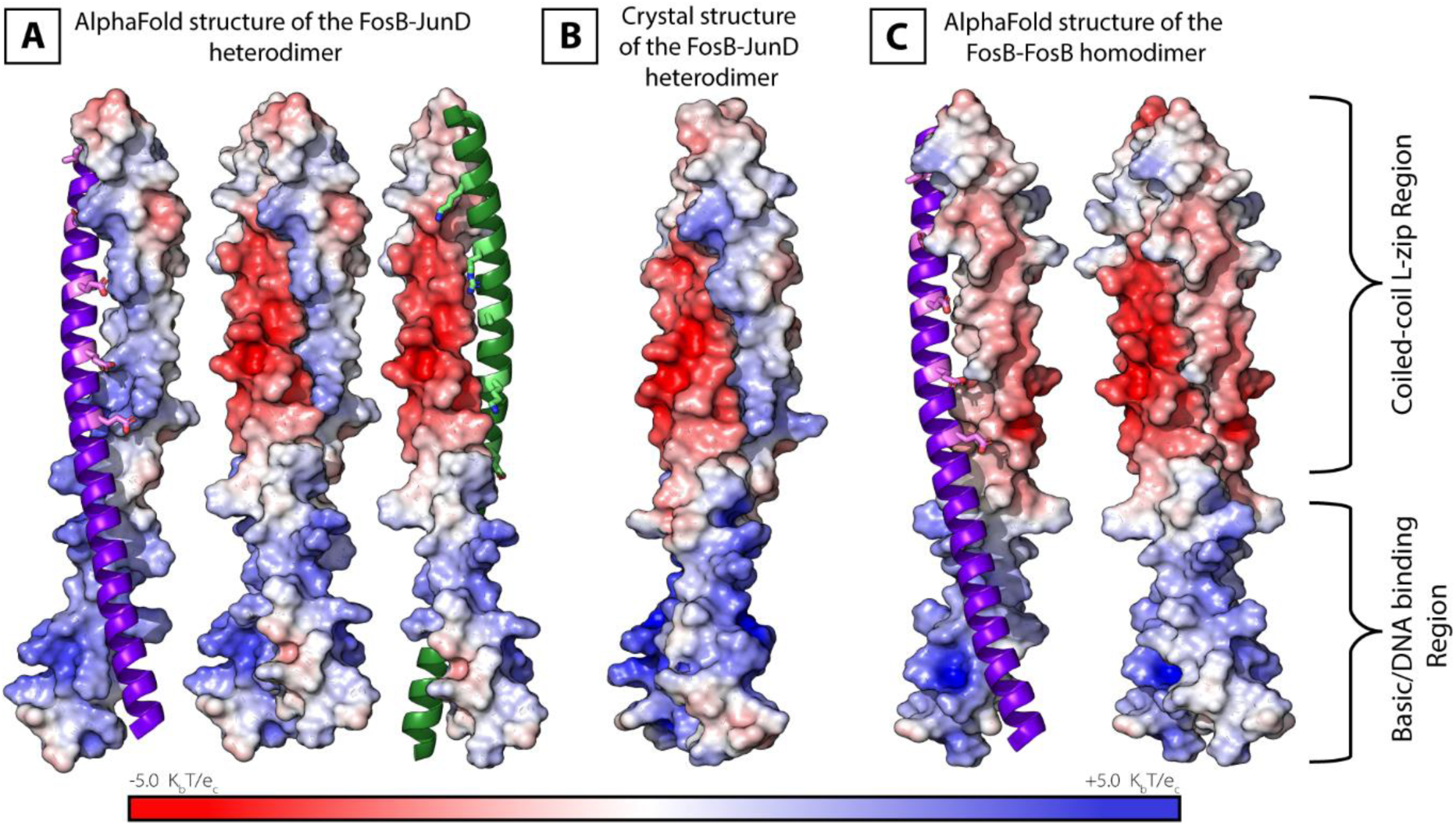
Surface potential of the structured coiled-coil region of: A- the AlphaFold-Multimer FosB-JunD heterodimer; B- the crystal structure FosB-JunD heterodimer, and C- the AlphaFold-Multimer FosB homodimer. Throughout: FosB shown in purple, and the g positioned residues are in violet and displayed as sticks. JunD is shown in green, and the e positioned residues are in lime green and shown as sticks.

The predicted FosB homodimer surface electrostatics told a different story. As can be seen in Figure 5 Panel C, the *e* residues of FosB created a less favourable surface potential for binding with the negatively charged FosB *g* residues. Consequently, the predicted structure of the FosB L-zip homodimer contained two negatively charged surfaces in close proximity. There was no indication in the pLDDT scores, the PAE scores or the ipTM scores (Full-length constructs), that the FosB homodimer would contain such an implausible dimer interface. When in fact, these clashing negative surface potentials are strong indicators that this dimer construct should not be able to form *in vivo*.

The electrostatic surface potential of the JunD homodimer was less clear. JunD had a neutral to basic overall surface potential for its coiled-coil region (See Supplementary Figure 6), without a clear pattern of opposing or repelling charges. It has been shown experimentally that JunD can homodimerize *in vitro* (Newman and Keating, 2003), and these results are consistent with that.

Thus far, we have shown that AlphaFold2 and AlphaFold-Multimer can recognise and accurately calculate the structures of L-zip dimers. We have also shown that AlphaFold-Multimer does not clearly differentiate between probable and improbable L-zip dimer interfaces, and therefore, produced high-confidence structures of the FosB homodimer, despite previous studies showing that FosB was unlikely to homodimerize (Newman and Keating, 2003; Vinson, Acharya and Taparowsky, 2006).

### Other human L-zip pairs show similar results to FosB and JunD

We wanted to investigate if the phenomenon of AlphaFold-Multimer producing high confidence and low PAE scoring L-zip dimers, even when the particular pair of proteins does not dimerise *in vivo*, was unique to Fos and Jun. To do this, we ran four more pairs of human proteins containing L-zip regions through AlphaFold-Multimer (Evans *et al*., 2022). All four protein pairs have been shown not to form stable dimers (Newman and Keating, 2003). As can be seen in Figure 6, all the different L-zip pairs that we ran were predicted to form L-zip dimers by AlphaFold-Multimer. They all had structured L-zip domains with high pLDDT scores and corresponding low PAE scores, indicating good AlphaFold-Multimer predictions that should be replicable *in vitro*. These results were all consistent with the previous Fos/Jun dimerisation case study. Looking at the AlphaFold-Multimer metrics alone, there is little to nothing to show that these L-zip pairs are in fact unstable dimer structures.

**Figure 6:**
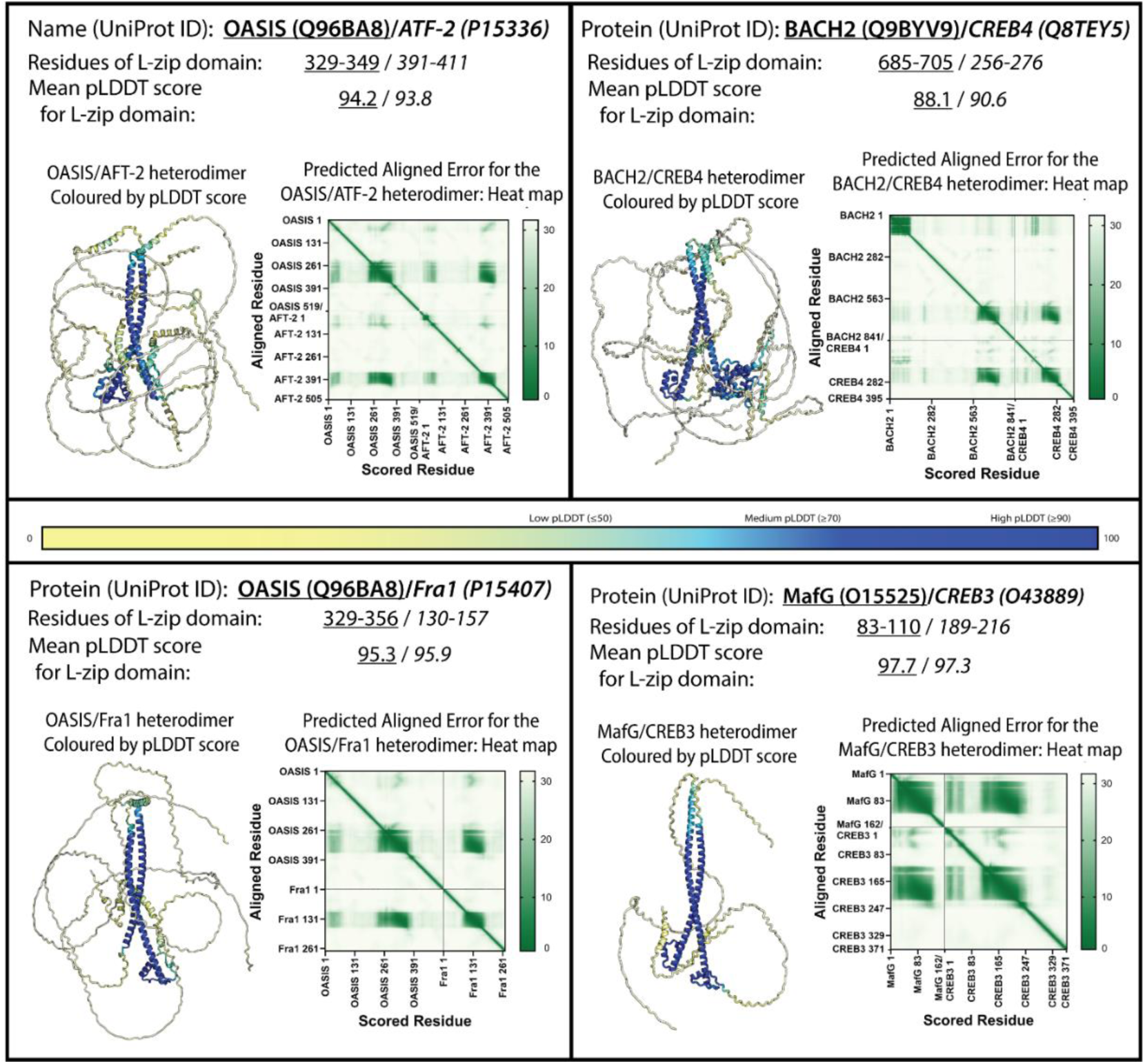
Four different human L-zip dimer constructs predicted by AlphaFold-Multimer (Evans et al., 2022). All four dimers have been experimentally demonstrated not to form in vitro (Newman and Keating, 2003).

### An energetically unfavourable synthetic peptide pair still forms a L-zip dimer

Next, we tested if AlphaFold-Multimer would form a L-Zip from a synthetic peptide, specifically engineered to be as energetically unfavourable as possible. Vinson et al., (2002; 2006) calculated the coupling energy (ΔΔΔG_int_) of common *g-e* pairs, relative to an Ala-Ala pair, and their data were used as a guide to design, *in silico,* two peptides with the highest possible coupling energy. Each peptide contained five heptad repeats, and the *e* and *g* positions were filled with repelling negatively charged residues. The first iteration of this peptide contained all Glu residues. (See Figure 7 panel B) (See supplementary Figure 1 for the monomer structures.)

**Figure 7:**
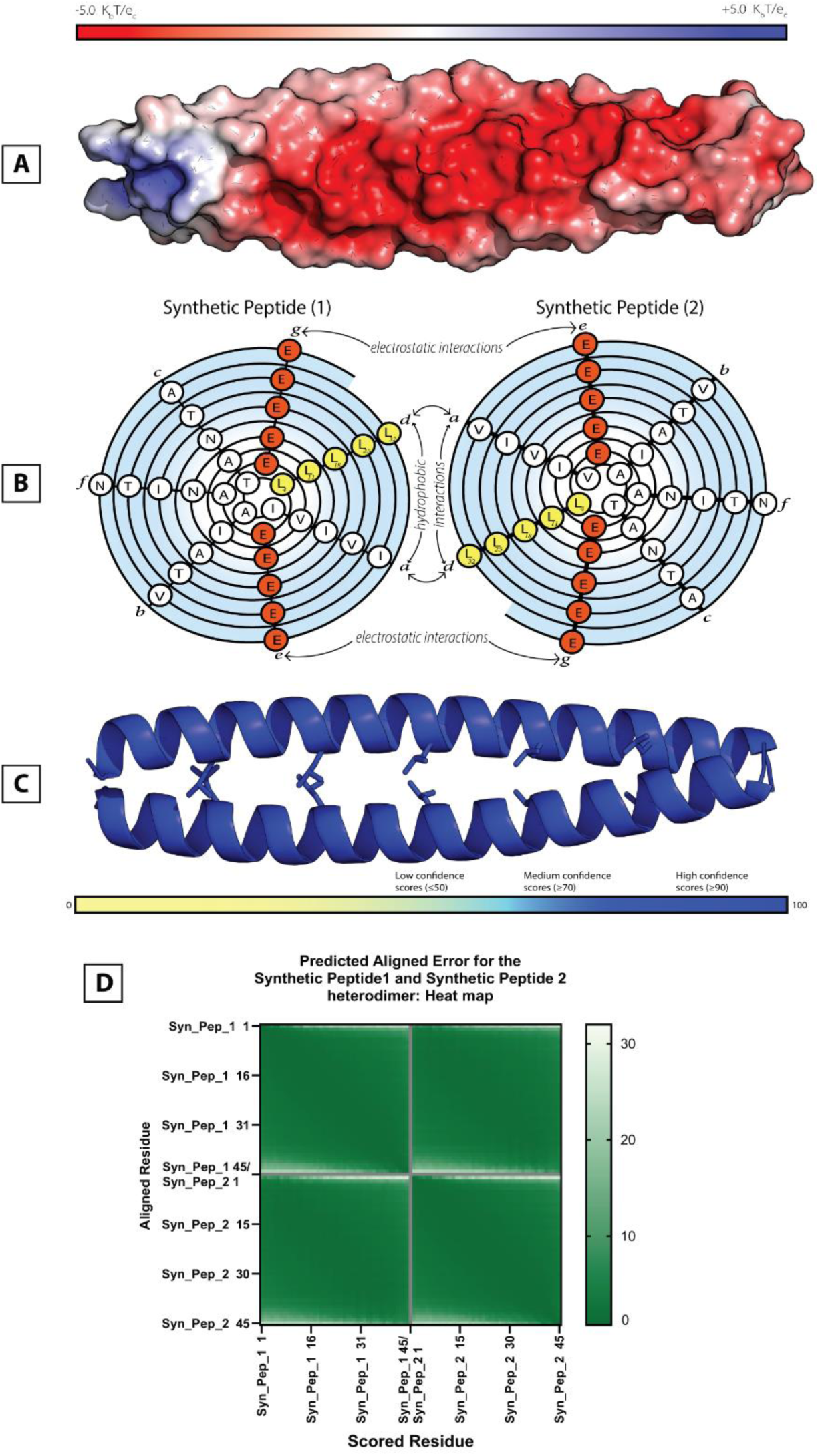
Surface potential, spiral diagram of the sequence, and structure of the synthetic L-zip peptide pair. A: Surface of the synthetic peptide coloured by surface electrostatic potential. B: Primary sequence in a spiral diagram, negatively charged residues are shown in red and Leu residues are shown in yellow. C: Predicted structure with Leu residues shown as sticks, coloured by confidence scores. D: The Predicted Alignment Error plot.

It should be noted that *de novo* protein design using AlphaFold is still difficult and an ongoing area of research (Goverde et al., 2023). The results from the following experiments show some of the problems associated with using AlphaFold2 and AlphaFold-Multimer to predict synthetic protein structures.

Despite the egregious clashing charges, AlphaFold-Multimer formed a L-Zip dimer from these synthetic peptides, with very high pLDDT scores (mean: 98.04) and very low PAE scores. (See Figure 7. Panel C, D.) An alignment between the synthetic peptide and the FosB-JunD crystal structure, showed that the dimer structure had been formed with coiled-coil L-zip geometry: RMSD: 0.984Å. (See Supplementary Figure 5.) As can be seen on Figure 3 Panel B, the ipTM scores for our synthetic peptide pair were also high, highest: 0.81, similar to the ipTM scores for the FosB/JunD heterodimer (L-zip region only). In this case, it appeared there was no way to distinguish the dimerization propensity of our synthetic peptide pair from that of the FosB-JunD dimer; both predicted structures had excellent confidence scores, but drastically different chances of forming *in vivo*.

### L-zip dimer prediction is dominated by leucine patterning and the commonness of *e* and *g* position residues, not electrostatics

As a final experiment, we wanted to test what would “break” the L-Zip dimer by using the synthetic peptide pair as a backbone and changing either the *e/g* residues, or the Leu (*d*) residues. Either all the *e/g* residues were changed to the same charged residue, or one, two or three Leu residues were changed to Ile residues. A predicted L-zip construct was recorded as “broken” when it included anti-parallel helices, parallel helices with incorrectly oriented Leu residues, or helices with very minimal interchain interaction. (See Table 2.)

**Table 2:**
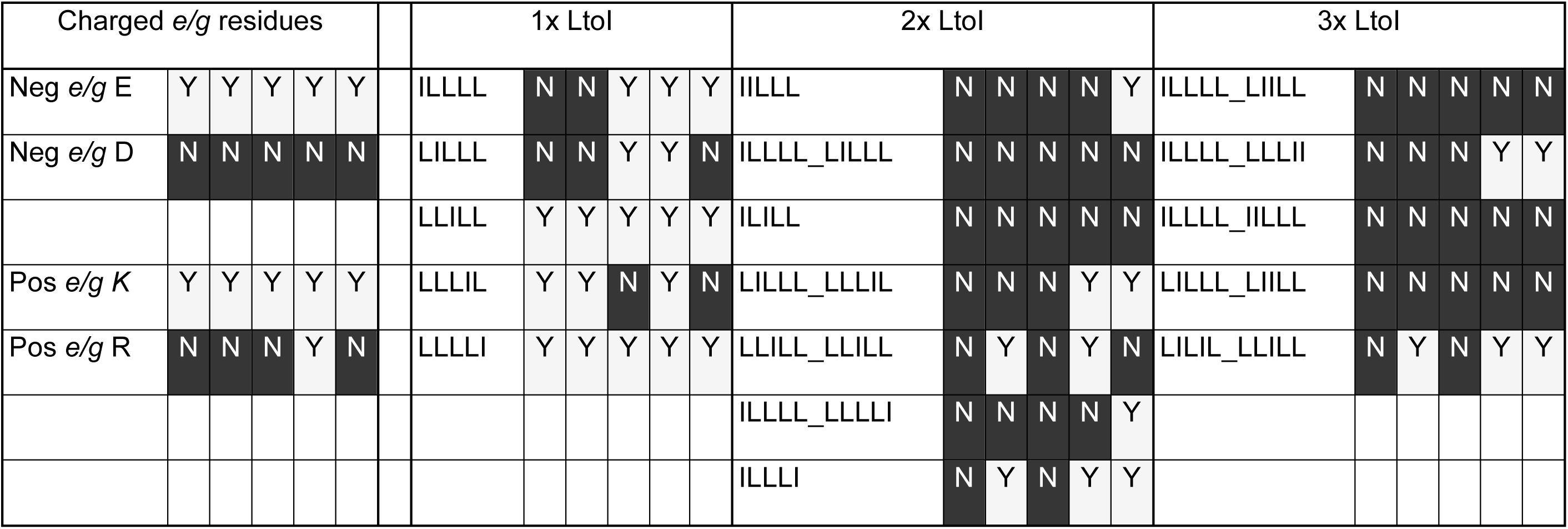
Results of synthetic L-zips in AlphaFold-Multimer. Any synthetic peptide pair that produced a dimeric coil-coil L-zip structure in AlphaFold-Multimer was marked as Y for Yes. Any peptide pair that produced any other structure in AlphaFold-Multimer was marked N for No. AlphaFold-Multimer was set to produce five ranked structures, each structure was assessed separately, producing five Y or N scores for each Synthetic peptide pair. (See Supplementary material for PDB files.)

Placing Glu and Lys residues in all the *e/g* positions still produced L-zips in all five AlphaFold-Multimer constructs. However, Asp and Arg *e/g* residues “broke” the L-zip, and caused AlphaFold-Multimer to produce alternative dimer interfaces. Sequence alignments of human L-zip proteins show that Arg and Asp are less commonly found in the *e/g* position than Lys or Glu (Vinson *et al*., 2002), and this could explain why AlphaFold-Multimer stopped recognising the L-zip heptad repeats when so many uncommon residues were included in the primary sequence. Changing one Leu to an Ile mostly maintained the L-zip construct, whereas, progressively mutating the Leu residues to Ile increased the number of “broken” L-zip constructs. (See Table 2.)

## Discussion

The neural network at the heart of AlphaFold2 and AlphaFold Multimer (Evoformer) builds and progressively refines a representation of the proximity of protein residues in the form of an evolving graphical network (Jumper *et al*., 2021; Evans *et al*., 2022). The evidence presented in this study demonstrates how AlphaFold2 cannot account for other effects (for example electrostatic) that can modify proximity. This is because the edges in the graphical network encode only the nodes that they join. Additionally, the relaxation algorithms utilized by AlphaFold2 and AlphaFold-Multimer (the Amber ff99SB force field) resolve steric clashes, but do not take into account destabilising electrostatic interactions (Hornak *et al*., 2006; Bouatta, Sorger and Al Quraishi, 2021; Jumper *et al*., 2021). L-zips are an excellent demonstration of these limitations: AlphaFold uses multiple sequence alignments to accurately find the L-zip regions, without recognising when clashing charges make the dimer structure improbable.

At the time of writing, AlphaFold3 is available to beta test and these limitations remain in this latest iteration of the software (Abramson *et al*., 2024).

The points raised in this study are important to consider when using AlphaFold to study any protein that forms homodimers or heterodimers as part of a combinatorial control network, or any proteins that form coiled-coil dimers. As a specific example, disruptions to the network of AP-1 transcription factors have been implicated in the development of bone, skin, liver and lung cancer, to name but a few (Eferl and Wagner, 2003). Assessing the different possible interactions within this nuanced network of transcription factors would be challenging using AlphaFold-Multimer. Similarly, to use AlphaFold2 and AlphaFold-Multimer to design artificial dimers, would require the use of additional data or software to evaluate binding partner specificity and affinity.

In the case of L-zips, previous experimental data can be used to evaluate the AlphaFold-Multimer structures. The coupling energies experimentally calculated by Vinson et al., (2002; 2006) can be used to estimate if a L-zip dimer will be stable. Unfortunately, Vinson et al., (2002; 2006) only looked at the most commonly found residues in these positions, therefore, it could not be used to accurately calculate the coupling energy of the FosB-JunD dimer, because JunD contains a relatively rare Thr as one of its *g* residues. However, previous experimentation like this can, and should, be used to assess the quality of AlphaFold structures. It should be emphasised that AlphaFold Multimer does not provide a theoretical K_D_ for predicted structures, or any other indication of binding strength.

Altogether, this study is an important example of an AlphaFold structure with high confidence scores but low accuracy. It illustrates AlphaFold’s strengths and its weaknesses, showing that AlphaFold structures must be interpreted critically, and with prior knowledge of the physical principles affecting protein 3D geometry.

## Methods

AlphaFold2 and AlphaFold multimer were accessed using the Galaxy Australia platform (Community, 2024), running AlphaFold v2.3.2. All AlphaFold computations were run using the default settings, as implemented by Galaxy Australia, with Amber99sb force field (Hornak *et al*., 2006) relaxation enabled. Five PDB protein structure files were produced, as per the default settings, for each AlphaFold computation. These PDB models were ranked 0 to 4, with 0 being the top ranked model, as calculated by Galaxy Australia. The model ranked 0 PDB, and all corresponding data files produced, were used for all the downstream analysis, unless otherwise stated. In all the AlphaFold computations discussed, a FASTA file containing the primary sequence of the protein/s was the only data input.

This study used full length sequences of FosB and JunD, sourced from UniProt (The UniProt Consortium, 2019), to ascertain if AlphaFold2 (Jumper *et al*., 2021; Evans *et al*., 2022) can recognise the L-zip motifs within the full sequences.

FosB (UniProt ID: P53539) and JunD (UniProt ID: P17535) were input into AlphaFold2 as monomers, homodimers, and a heterodimer. All other protein primary sequences discussed, apart from the synthetic peptide pairs, were sourced from UniProt (The UniProt Consortium, 2019), with the UniProt ID numbers listed in the text.

In order to analyse the per-residue local distance difference test (pLDDT) scores, they were extracted from the B-factor column of the PDB files produced in our AlphaFold computations.

The structure alignments were calculated with the Pymol Alignment plugin using the default settings (Schrodinger LLC, 2015). (Pymol Command: extra_fit *x*, y, \ method=align, \ cycles=5, \ cutoff=2.0, \ mobile_state=-1, \ target_state=-1.) For each structural alignment discussed, the AlphaFold predicted structure was aligned to the target crystal structure of the FosB-JunD heterodimer (PDB ID: 5VPF) (Yin, 2019). The L-zip domain from the crystal structure was isolated, FosB(T180-V214) and JunD(I293-K327), and the corresponding residues from the AlphaFold predicted structures were also isolated and aligned to the target. For the synthetic peptide alignments, residues 6-40 of synthetic peptides 1 and 2 were isolated and aligned to the L-zip domain, FosB(T180-V214) and JunD(I293-K327), of the above crystal structure target. Isolating the L-zip domains removed undue influence from the unstructured/highly variable regions of the FosB/JunD predicted AlphaFold structures and ensured consistency across the different structural alignments.

The surface electrostatics were calculated using the Pymol APBS electrostatics plugin using the default settings (Schrodinger LLC, 2015; Jurrus *et al*., 2018).

When required, .pkl files were produced (one per ranked PDB model), imported into the molecular graphics software UCSF ChimeraX (Meng *et al*., 2023), and used to visualise the predicted alignment error (PAE) scored pseudo-bonds.

When required, predicted-alignment error (PAE) matrix .csv files and .txt files containing the ipTM scores were produced, and imported into GraphPad Prism Version 10.2.0 (GraphPad Software), and used to make heat maps of the PAE scores and other figures.

## Supporting information

Synthetic Leucine zipper construct PDB files

## Acknowledgments

This work was supported by a National Health and Medical Research Council Ideas Grant (GNT2013215; D.A.J., T.B.) and Wellcome Trust Collaborator Award (214344/Z/18/Z; D.A.J.). D.A.J. was supported by a UNSW Scientia Fellowship. We also acknowledge the use of the Structural Biology Facility in the Mark Wainwright Analytical Centre – UNSW, funded in part by the Australian Research Council Linkage Infrastructure, Equipment and Facilities Grant: ARC LIEF 190100165. We acknowledge the Bedegal people of the Eora nation, the traditional custodians of the land upon which this research took place. We’d like to thank Professor Peter Mitic, University College London Department of Computer Science, for help with statistical analysis.

## Notes

### Competing Interest Statement

The authors have declared no competing interest.

