## Supplementary figures 1-6 for "Assessing the Validity of Leucine Zipper Constructs Predicted in AlphaFold2"

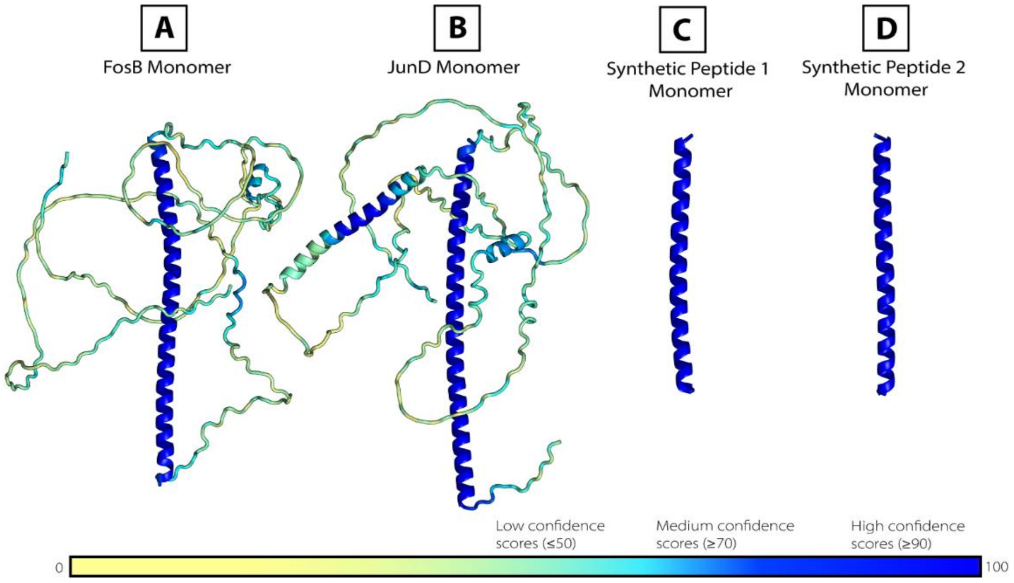


*Supplementary Figure 1: AlphaFold2 structures of the monomers of: A- FosB; B- JunD; C- Synthetic Peptide 1; D- Synthetic Peptide 2. All structures have been coloured by confidence score, Dark blue shows high confidence (>90), pale yellow shows low confidence (<50).*

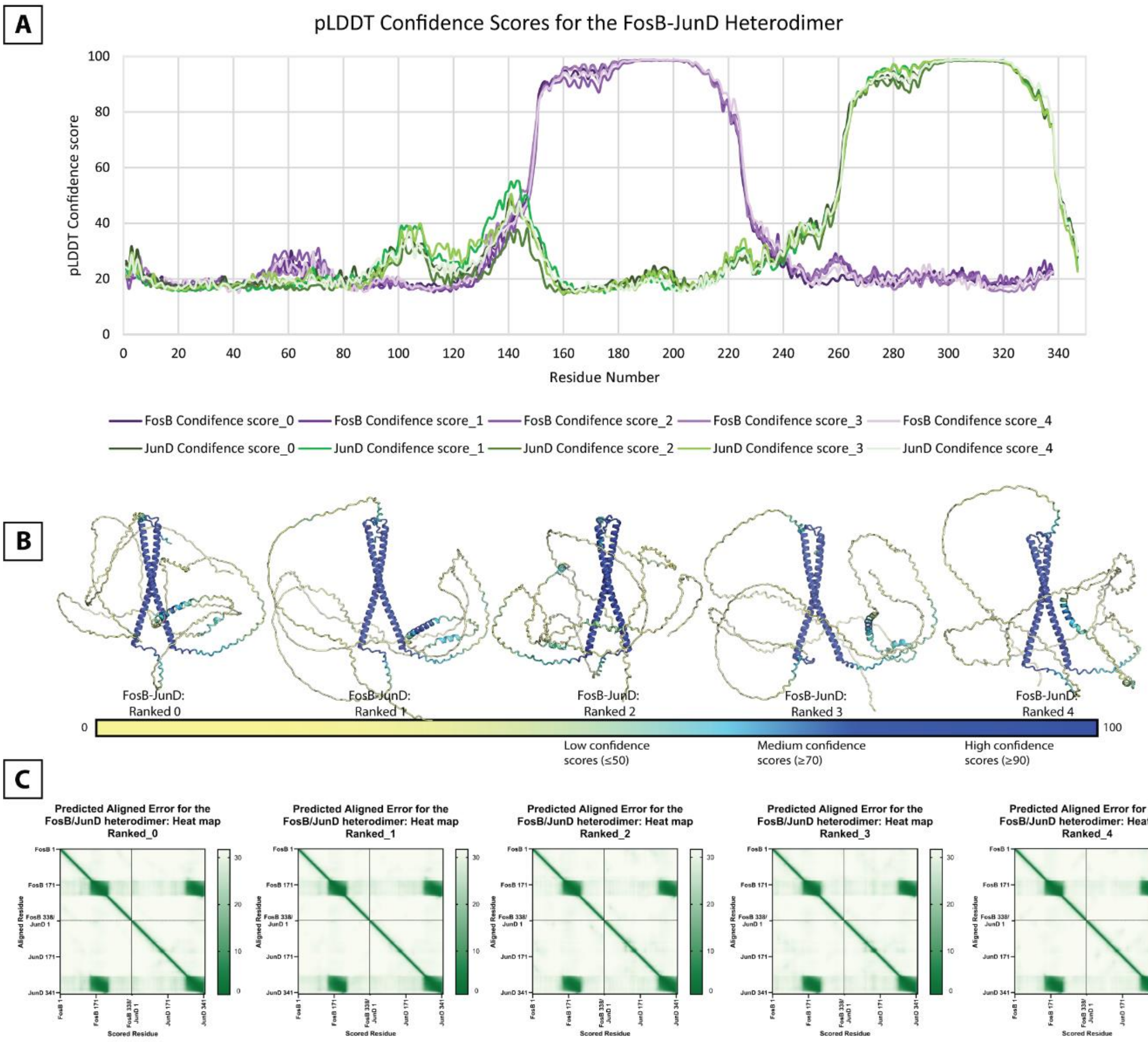


*Supplementary Figure 2: Confidence scores for the FosB-JunD Heterodimer and the structures for each of the ranked constructs that AlphaFold2 produced, showing high similarity between all five constructs. A: Graph of the pLDDT scores for ranked structures 0-4, coloured by chain. B: FosB-JunD Heterodimer structures are coloured by confidence score. C: PAE heatmap plots for ranked structures 0-4.*

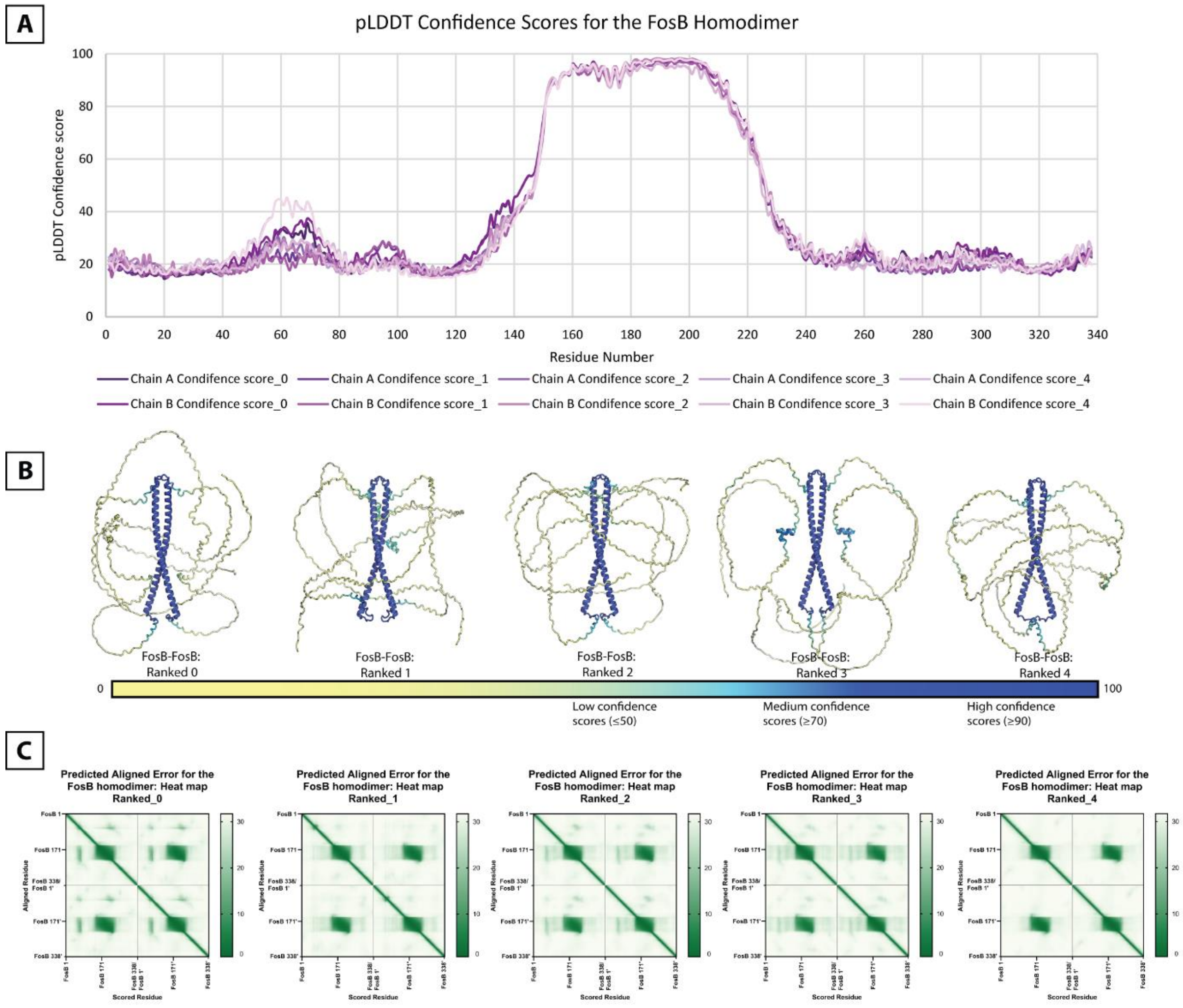


*Supplementary Figure 3: Confidence scores for the FosB-FosB Homodimer, and the structures for each of the ranked constructs that AlphaFold2 produced, showing high similarity between all five constructs. A: Graph of the pLDDT scores for ranked structures 0-4, coloured by chain. B: FosB-FosB Homodimer structures are coloured by confidence score. C: PAE heatmap plots for ranked structures 0-4.*

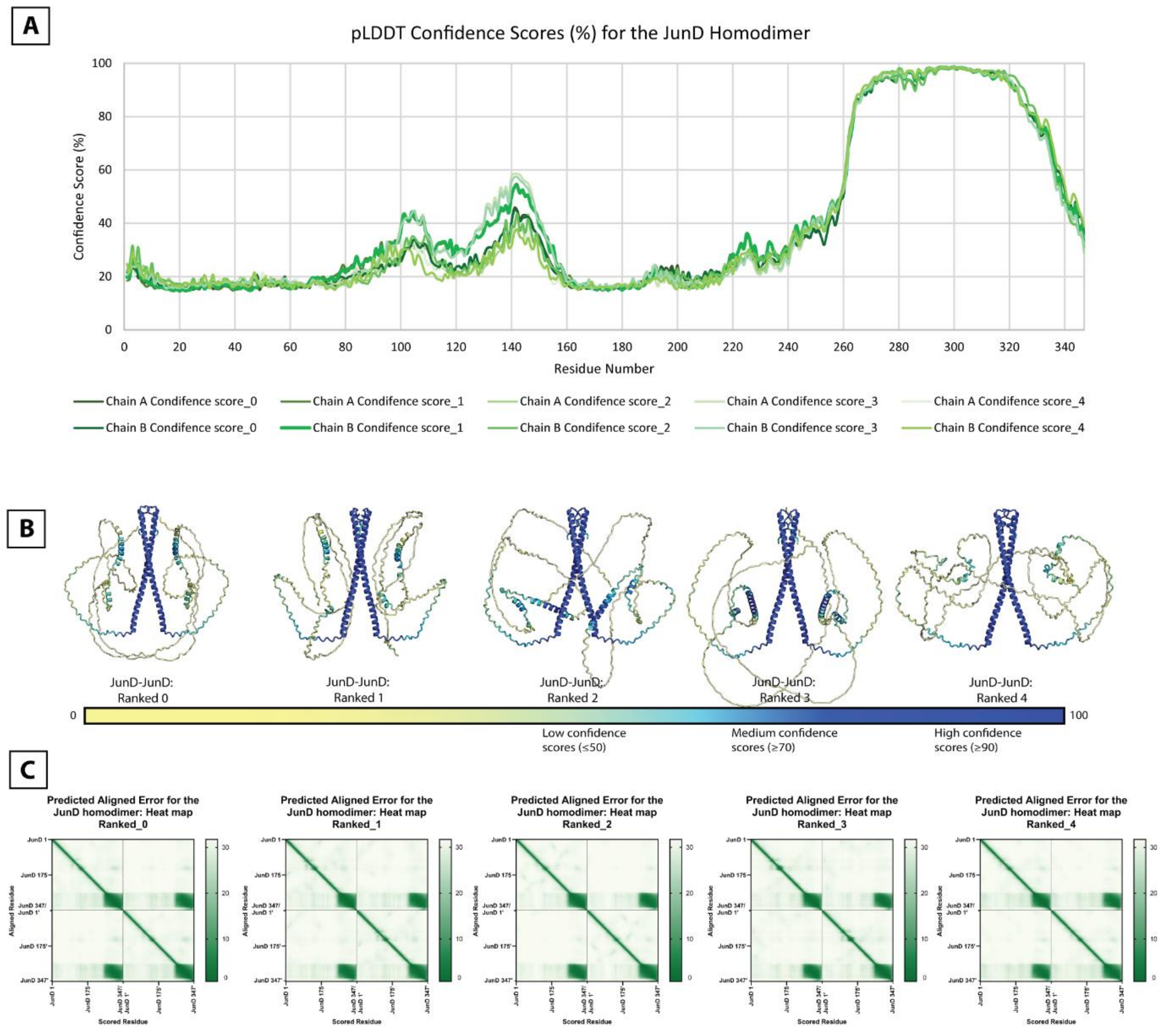


*Supplementary Figure 4: Confidence scores for the JunD-JunD Homodimer, and the structures for each of the ranked constructs that AlphaFold2 produced, showing high similarity between all five constructs. A: Graph of the pLDDT scores for ranked structures 0-4, coloured by chain. B: JunD-JunD Homodimer structures are coloured by confidence score. C: PAE heatmap plots for ranked structures 0-4.*

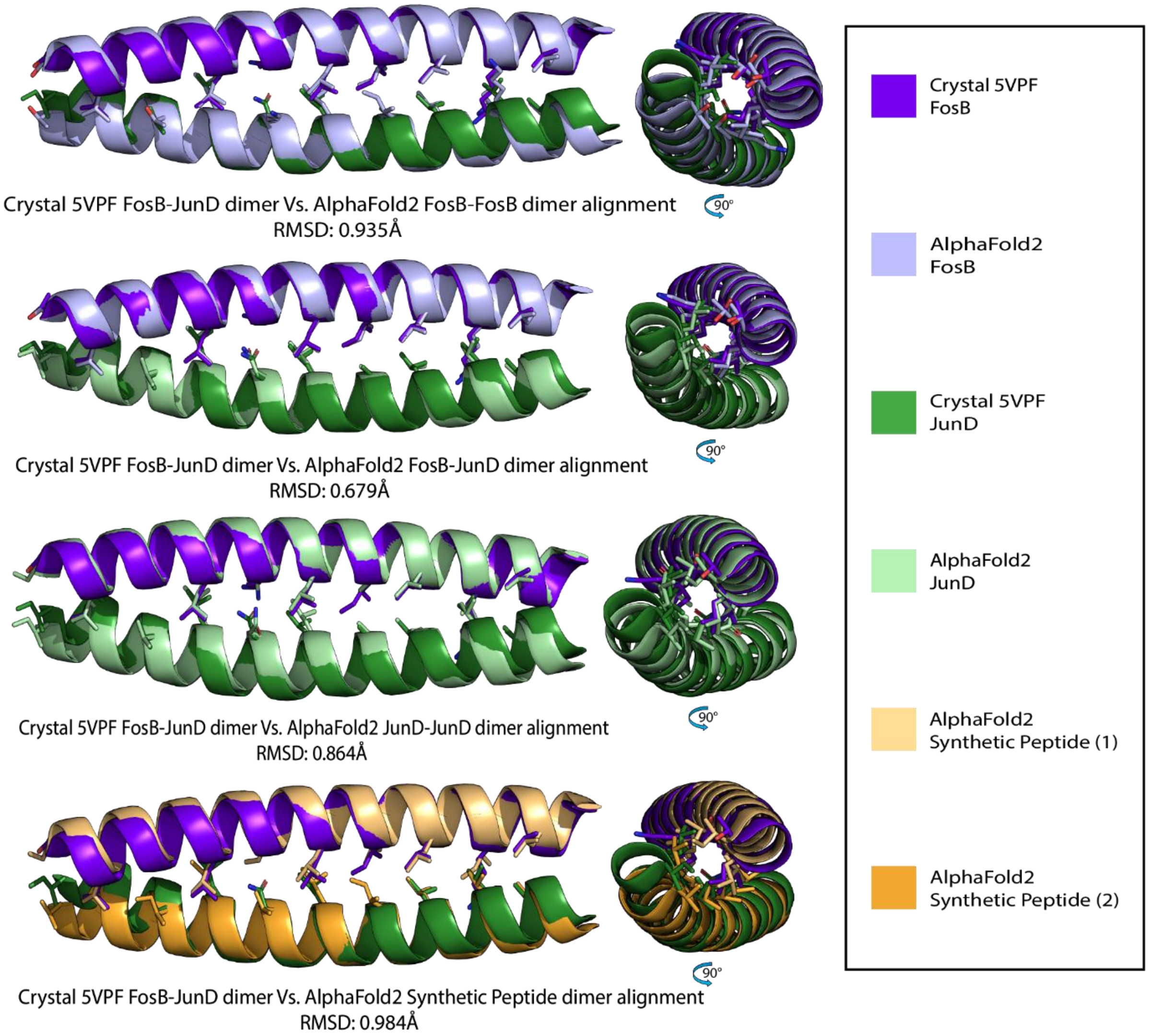


*Supplementary Figure 5: Alignments between the crystal structure of the FosB-JunD heterodimer (5VPF) and different AlphaFold2 structures. This demonstrates the close structural alignments between the AlphaFold2 L-zip structures and the experimental L-zip structure.*

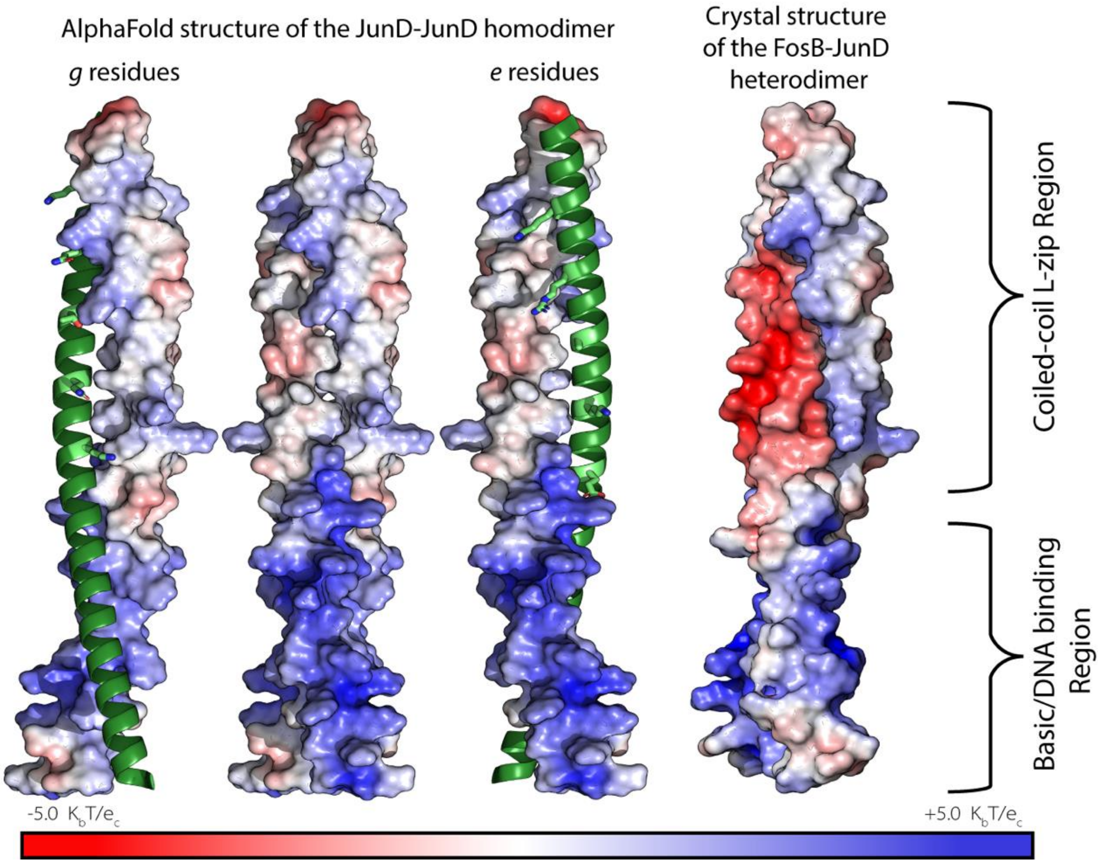


*Supplementary Figure 6: Surface potential of the structured coiled-coil region of the AlphaFold2 JunD homodimer and the crystal structure FosB-JunD heterodimer. The cartoon representation of JunD is shown in green, and the e or g positioned residues are shown in lime green and as sticks.*

Supplementary material: .zip file of all the PDB files for the synthetic peptide pairs structures produced in AlphaFold-Multimer.

File name: Synthetic Leucine zipper construct PDB files.zip
